# Effects of Exercise Frequency on Aortic Calcification in Hyperlipidemic Mice

**DOI:** 10.1101/2024.04.03.588017

**Authors:** Nora Safvati, Sophia Kalanski, Andy Hon, Stuti Pradhan, Mimi Lu, Linda L. Demer, Yin Tintut

**Author notes:** To whom correspondence should be addressed Yin Tintut, Ph.D., The David Geffen School of Medicine, University of California, Los Angeles, Center for the Health Sciences A2-237, 10833 Le Conte Ave, Los Angeles, CA. 90095-1679.

## Abstract

Cardiovascular disease risk is associated with coronary artery calcification and is mitigated by regular exercise. However, the optimal regimen (activity level and frequency) for benefitting cardiovascular health is not clear. The surface area of calcium deposits, in addition to the quantity of calcium mineral, is a key factor in the risk of plaque rupture in humans due to compliance mismatch of edges with surrounding distensible tissue resulting in debonding. We previously found that aortic PET tracer uptake, a marker of mineral surface area, was reduced in hyperlipidemic (*Apoe*^*-/-*^) mice that underwent treadmill exercise, especially those in the low-speed (12.5 meters/min) regimen. To test the optimal exercise frequency on cardiovascular health, hyperlipidemic mice with baseline aortic calcification were subjected to none, 3- or 5-day/week treadmill regimen (12.5 meters/min for 30 min/day) for 5 weeks. MicroPET/microCT imaging and echocardiography were performed before and after the 5-week study. Results show that aortic calcification significantly progressed in all 3 groups, and ^18^F-NaF uptake increased in the control and 3-day groups, but not in the 5-day group. As for cardiac function, left ventricular (LV) systolic function was not affected but LV mass and wall thickness decreased in the 5-day group, suggesting cardiac remodeling. Skeletal bone density assessed at lumbar vertebrae shows that bone density decreased in all 3 groups. These findings suggest that, in mice with underlying calcific atherosclerosis, low-speed exercise and a frequency of 5 days/week is optimal for benefitting cardiovascular health.

## Introduction

Physical activity is associated with lowering risk for cardiovascular events and mortality [1]. However, the optimal regimen (activity level and frequency) for benefitting cardiovascular health is not clear. A U-shaped association has been reported for the physical activity level and cardiovascular events [2]. A recent study shows that exercise intensity, but not exercise volume, is associated with changes in coronary artery calcification (CAC) [3]. Prevalence of cardiovascular disease and risk factors were lower with low and moderate volumes of exercise but not further benefited by higher doses of exercise [4].

The amount of cardiovascular calcium content, especially CAC, is known to associate with risk for adverse cardiovascular events and mortality [5, 6]. Interestingly, a paradoxical relationship is found in elite athletes where, compared with less active controls matched for conventional risk factors, veteran male elite athletes have greater prevalence and severity of CAC [7-10], although they are not generally associated with greater cardiovascular risk. In fact, cardiovascular outcomes of elite athletes appear to be better than the general population [11]. One possibility is that elite athletes have less mineral surface area exposed to distensible tissue, since the morphology of their plaques are composed more of calcium than lipids [12]. By theoretical analyses, rupture risk is the greatest in the area between mineral edges and surrounding distensible tissue [13], suggesting that qualitative features, such as mineral surface area, may also influence plaque stability. Our recent studies in female hyperlipidemic mice demonstrate that aortic ^18^F-NaF PET uptake, a marker of surface area, was reduced in mice that underwent a progressive exercise regimen [14] as well as a constant low-speed exercise regimen [15]. Thus, in this study, we tested the optimal exercise frequency on a qualitative feature of aortic calcification.

## Materials and Methods

### Animals and treadmill exercise regimen

Experimental protocols were reviewed and approved by the Institutional Animal Care and Use Committee of the University of California, Los Angeles.

Given that our prior study [15] showed exercise affected aortic ^18^F-NaF PET uptake only in female mice, and not in male mice, we used only female mice in this study. *Apoe*-null hyperlipidemic mice (n = 44, ∼ 8-month-old female retired breeders on C57BL/6 background, Jackson Laboratory) were placed on a “Western” diet (21% fat and 0.2% cholesterol, Envigo) to induce baseline aortic calcification for approximately 2 months. Prior to the start of the experiment, the mice were acclimated to the treadmill experience (Columbus Instruments Exer-3/6 Animal Treadmill Rodent 6-Lane) for 3 – 4 days followed by a one-time, 10 min, test session for running capacity at 12.5 m/min. The 3 mice that did not tolerate the test were excluded from the subsequent experiment. After undergoing microCT/PET imaging and echocardiography, the mice were divided into 3 groups (13 mice/group) - control (no treadmill), 3-day (3 days/week), or 5-day (5 days/week) - and subjected to treadmill running for 5 weeks. The speed and duration of each treadmill run (0° slope, no electric shock stimulation) for 3-day and 5-day groups was kept constant at 12.5 m/min and 30 minutes, respectively, for the entire 5-week study. To reduce any potential confounding effects of the Western diet, all the mice was placed on the standard diet for the duration of the exercise regimen. Two mice in the 3-day group were excluded due to premature death.

### Serial in vivo ^18^F-NaF microPET/microCT imaging and analysis

Fused ^18^F-NaF microPET/microCT images were acquired before the exercise regimen and again at Week 5 at the Preclinical Imaging Facility of the Crump Institute for Molecular Imaging at the California NanoSystems Institute at UCLA. The imaging and analysis protocols were described previously [16]. Briefly, mice were injected with ∼90 µCi ^18^F-NaF via the tail vein. One-hour post-injection, the mice were anesthetized and imaged in the microPET/microCT scanner (GNEXT).

Images were analyzed using AMIDE software [17]. CT quantification for aortic calcification was performed by three-dimensional isocontour (automated edge detection) regions of interest (ROIs) with a minimum threshold of 250 Hounsfield units (HU). PET quantification for aortic calcification was performed from an isolated volumetric ROI encompassing parts of cardiac and aortic regions with a minimum ^18^F-NaF isocontour threshold of 2% injected dose per cubic centimeter (%ID/cc). The mean threshold of background ^18^F-NaF uptake, measured at the cardiac silhouette of four mice, was 0.8 %ID/cc. The CT mineral content and the total PET uptake were each determined from values of mean density and volume (vHU or PET uptake). CT quantification for the lumbar bone density was performed by geometric box ROI analysis encompassing the L3 vertebra with a threshold of 1000 HU, and the mean density (HU) was used for the comparative analysis.

### Echocardiography

Mice were anesthetized (3.0% isoflurane for initiation and 1.5–2.0% isoflurane for maintenance delivered via nose cone), and M-mode and tissue Doppler echocardiography of the left ventricle were performed using a VisualSonics Vevo 3100 equipped with a 30-MHz linear transducer.

### Statistical analysis

Values are expressed as mean ± SEM. Statistical analysis was performed with Prism software (GraphPad, v. 9.4.1). Paired analysis was used for comparison of data of the same mouse between different time points. For comparisons among multiple groups with repeated measures, two-way ANOVA followed by Holm-Sidak post-hoc analysis was employed. All tests were 2-sided, and a *p* value ≤ 0.05 was considered statistically significant.

## Results

### Effects of exercise frequency on progression of aortic calcification

To assess aortic calcium content and mineral surface area, microCT/microPET imaging was performed on all mice at 2 time points: before and after the 5-week treadmill regimen (**Fig. 1A**). MicroCT analysis showed that aortic calcification progressed significantly in all 3 groups over the 5-week period (**Fig. 1B**). The progression was not significantly different among the 3 groups. However, the microPET analysis showed that PET-tracer (^18^F-NaF) uptake increased significantly over the study period in the control and 3-day group, but the uptake did not increase in the 5-day group (**Fig. 1C**).

**Figure 1.**
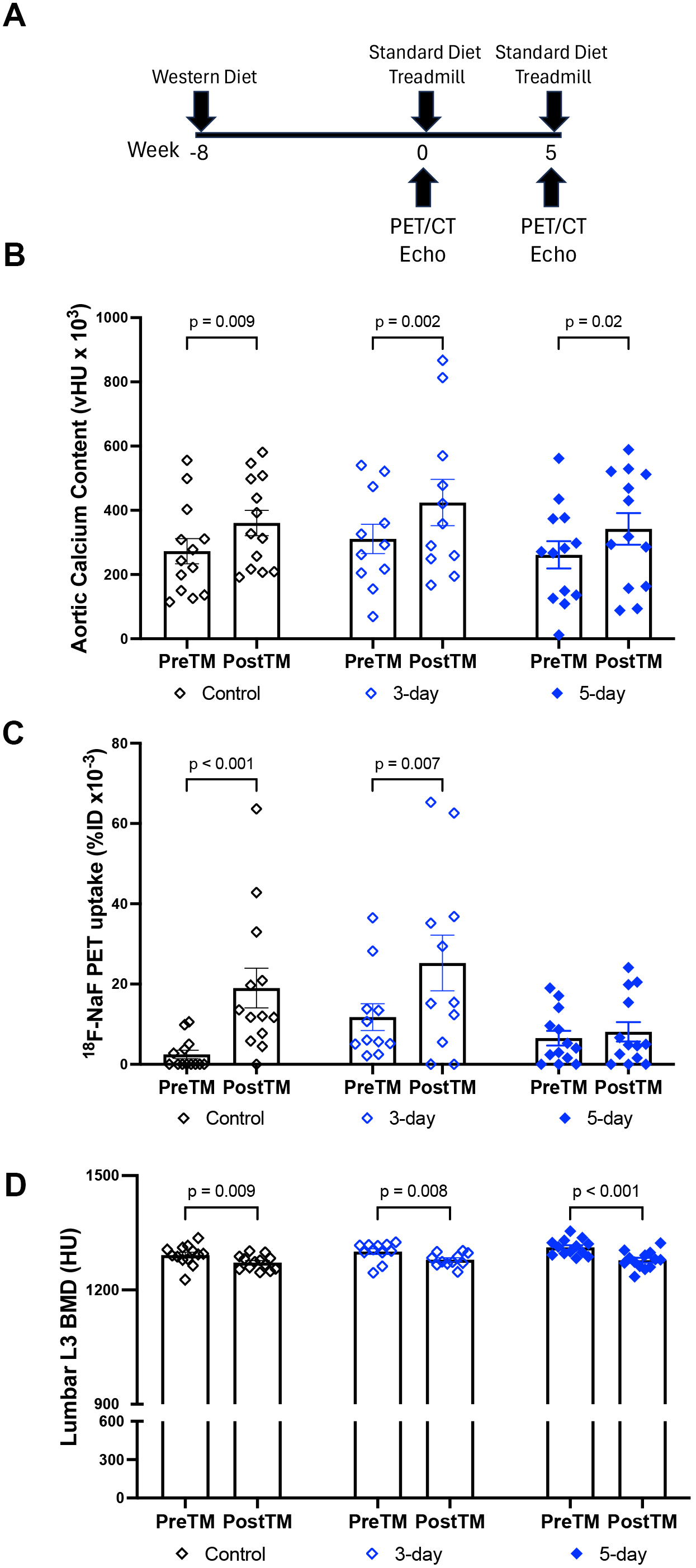
Effects of exercise frequency on aortic calcification and skeletal bone density. **(A)** Schematic diagram of experimental design. **(B)** Aortic calcium content assessed by microCT imaging. **(C)** Aortic ^**18**^ F-NaF uptake assessed by microPET imaging. **(D)** Skeletal bone density at lumbar vertebrae 3 assessed by microCT imaging.

### Effects of exercise frequency on cardiac structure and function

Echocardiography was performed at Weeks 0 and 5. As shown in **Table I**, left ventricular mass and wall thickness were significantly reduced in mice in the 5-day group compared with control and 3-day groups. Cardiac function was not significantly altered in all 3 groups.

### Effects of exercise frequency on skeletal bone mineral density

Bone density at lumbar vertebrae (L3) was assessed by microCT. As shown in **Fig. 1D**, vertebral bone mineral density decreased in all three groups.

## Discussion

Regular physical activity lowers cardiovascular risk [2]. Interestingly, prevalence of cardiovascular disease is reported to be lower with mild and moderate doses of lifelong exercise, but not further benefited by high doses of exercise [4]. We previously found that a low-intensity treadmill regimen (30 min/day for 5 days/week) decreased PET tracer uptake compared with controls or high-intensity regimen in female hyperlipidemic mice but not in male mice [15]. In this study, we tested the effects of exercise frequency on qualitative features of aortic calcification and quantitative cardiovascular measures in hyperlipidemic mice with underlying calcific atherosclerosis.

The surface area of calcium deposits, in addition to the quantity of calcium mineral, is a key factor in assessing risk of plaque rupture due to compliance mismatch of calcium (hydroxyapatite) mineral edges with surrounding distensible tissue. The mechanical mismatch leads to increased rupture (von Mises) stress, which, when it exceeds the strength of the soft tissue, results in separation at the surface of the deposit known as debonding [13, 18-20]. Irkle and colleagues elegantly showed that ^18^F-NaF uptake occurs at the surface of calcium deposits in human carotid arteries [21], which is consistent with the known adsorption and covalent binding of fluoride to hydroxyapatite mineral to form fluoroapatite [22]. In this way, the amount of ^18^F-NaF tracer uptake within plaque is a marker for the mineral surface area. Therefore, in this study, we used ^18^F-NaF PET imaging to quantify mineral surface area. Results showed that aortic ^18^F-NaF tracer uptake was significantly increased in the control and 3-day groups, but not in the 5-day group, suggesting that a 5-day/wk exercise regimen is sufficient and a frequency of more than 3 days/wk is needed to prevent progression of the vulnerable calcium deposit surface area. As for progression of aortic calcium content, as we have observed previously [14, 15], aortic calcification progressed significantly in all 3 groups, and exercise did not significantly affect progression over that of controls.

By echocardiography, cardiac function was not significantly changed with exercise in any of the 3 groups. In contrast, cardiac structure was affected by exercise: in mice on the 5-day exercise regimen, LV mass was significantly reduced as was wall thickness. This cardiac remodeling is consistent with changes seen in human subjects with cardiovascular dysfunction who underwent endurance exercise regimens compared with sedentary controls. In one study of a small number of hypertensive adults [23], 3 to 9 months of endurance exercise led to a significant 12% decrease in wall thickness and a significant reduction in LV mass index, whereas sedentary controls experienced no change or a slight increase in those parameters in controls. In the exercising subjects, the ratio of ventricular wall thickness to radius also declined by 15%; in contrast, there was no significant change in controls. In another study of middle-aged hypertensive adults undergoing endurance exercise training, a decrease in LV mass of borderline statistical significance was observed (9.4%; p = 0.056) [24]. Reduction of LV wall thickness was also reported in healthy postmenopausal women who underwent 4 months of walking training, and the reduction was correlated with leptin levels [25]. Thus, one possible mechanism of this unexpected finding is that the previous dietary hyperlipidemia superimposed on genetic hyperlipidemia resulted in cardiometabolic dysfunction, including both calcific atherosclerosis and concentric left ventricular hypertrophy, and that the ventricular hypertrophy regressed with exercise as observed in human subjects.

With respect to exercise effects on the skeleton, bone density decreased in the control and 3-day groups. Bone density also decreased in mice on the 5-day regimen, consistent with our previous study [15]. This is most likely as a result of hyperlipidemia, which we found previously to reduce skeletal bone density as well as anabolic effects of parathyroid hormone [26-29]. Another potential mechanism would be the endocrinological effects of exercise. For instance, in women, intense cross-country running is associated with amenorrhea and low bone mass [30].

Collectively, our present and prior findings in mice with underlying calcific atherosclerosis suggest that, while aortic calcification progresses over time irrespective of exercise, plaque vulnerability and left ventricular hypertrophy may regress in response to a low-speed treadmill exercise regimen on a 5-day/week schedule. These findings may provide a valuable context for exercise recommendation in subjects with underlying calcific atherosclerosis.

## Acknowledgments

The microPET/microCT imaging was performed at the Preclinical Imaging Facility of the Crump Institute for Molecular Imaging at the California NanoSystems Institute at UCLA, and we thank Dr. S. Xu and M. Tamboline for their assistance.

## Effects of Treadmill Frequency on Echocardiographic Parameters

| Parameters | Control (n = 12) |  | 3-Day (n = 11) |  | 5-Day (n = 12) |  |
| --- | --- | --- | --- | --- | --- | --- |
|  | Week 0 | Week 5 | Week 0 | Week 5 | Week 0 | Week 5 |
|  | Mean ± SEM | Mean ± SEM | Mean ± SEM | Mean ± SEM | Mean ± SEM | Mean ± SEM |
| Ejection Fraction (%) | 57 ± 3 | 56 ± 2 | 56 ± 4 | 59 ± 3 | 58 ± 3 | 59 ± 2 |
| Fractional Shortening (%) | 30 ± 2 | 29 ± 1 | 29 ± 3 | 31 ± 2 | 30 ± 2 | 31 ± 2 |
| Diameter, s (mm) | 2.5 ± 0.1 | 2.6 ± 0.1 | 2.4 ± 0.1 | 2.4 ± 0.2 | 2.5 ± 0.1 | 2.3 ± 0.1 |
| Diameter, d (mm) | 3.5 ± 0.1 | 3.7 ± 0.1 | 3.4 ± 0.1 | 3.4 ± 0.2 | 3.5 ± 0.1 | 3.3 ± 0.1 |
| LV Mass (corrected, mg) | 98 ± 4 | 99 ± 7 | 96 ± 6 | 86 ± 5 | 102 ± 3 | 81 ± 3<sup>a,b</sup> |
| LVAW,s (mm) | 1.5 ± 0.1 | 1.5 ± 0.1 | 1.4 ± 0.1 | 1.4 ± 0.1 | 1.6 ± 0.1 | 1.4 ± 0.1<sup>c</sup> |
| LVAW,d (mm) | 1.07 ± 0.04 | 1.03 ± 0.06 | 1.05 ± 0.07 | 0.99 ± 0.05 | 1.08 ± 0.05 | 0.94 ± 0.05<sup>c</sup> |
| LVPW,s (mm) | 1.13 ± 0.04 | 1.10 ± 0.04 | 1.22 ± 0.05 | 1.17 ± 0.05 | 1.27 ± 0.08 | 1.11 ± 0.05<sup>d</sup> |
| LVPW,d (mm) | 0.81 ± 0.03 | 0.77 ± 0.03 | 0.88 ± 0.03 | 0.82 ± 0.05 | 0.97 ± 0.08 | 0.79 ± 0.04<sup>e</sup> |

<sup>a</sup>p = 0.01 vs. Control (Week 5); <sup>b</sup>p < 0.001 vs. 5-Day (Week 0); <sup>c</sup>p = 0.05 vs. 5-Day (Week 0); <sup>d</sup>p = 0.02 vs. 5-Day (Week 0); <sup>e</sup>p = 0.001 vs. 5-Day (Week 0)
Diameter, systole (Diameter, s); diameter, diastole (Diameter, d); left ventricular mass (LV Mass); LV anterior wall, systole (LVAW;s); LV anterior wall, diastole (LVAW;d); LV posterior wall, systole (LVPW;s); LV posterior wall, diastole (LVPW;d)

